# Glucagon-like peptide 1 receptor-mediated stimulation of a GABAergic projection from the bed nucleus of the stria terminalis to the hypothalamic paraventricular nucleus

**DOI:** 10.1101/2021.03.04.433953

**Authors:** Nadya Povysheva, Huiyuan Zheng, Linda Rinaman

**Author notes:** Corresponding Author: Linda Rinaman, Ph.D., Professor of Psychology, Program in Neuroscience, Florida State University, 1107 W Call Street, Tallahassee, FL 32306, Voice. **Competing Interests Statement:** The authors have no competing interests to declare.

## Abstract

We previously reported that GABAergic neurons within the ventral anterior lateral bed nucleus of the stria terminalis (alBST) express glucagon-like peptide 1 receptor (GLP1R) in rats, and that virally-mediated “knock-down” of GLP1R expression in the alBST prolongs the hypothalamic-pituitary-adrenal axis response to acute stress. Given other evidence that a GABAergic projection pathway from ventral alBST serves to limit stress-induced activation of the HPA axis, we hypothesized that GLP1 signaling promotes activation of GABAergic ventral alBST neurons that project directly to the paraventricular nucleus of the hypothalamus (PVN). After PVN microinjection of fluorescent retrograde tracer followed by preparation of *ex vivo* rat brain slices, whole-cell patch clamp recordings were made in identified PVN-projecting neurons within the ventral alBST. Bath application of Exendin-4 (a specific GLP1R agonist) indirectly depolarized PVN-projecting neurons in the ventral alBST and adjacent hypothalamic parastrial nucleus (PS) via circuit-mediated effects that increased excitatory synaptic inputs and decreased inhibitory synaptic inputs to the PVN-projecting neurons; these effects were occluded by prior bath application of a GLP1R antagonist. Additional retrograde tracing experiments combined with *in situ* hybridization confirmed that PVN-projecting neurons within the ventral alBST/PS are GABAergic, and do not express GLP1R mRNA. Conversely, GLP1 mRNA is expressed by a subset of GABAergic neurons within the oval subnucleus of the dorsal alBST that project into the ventral alBST. Our novel findings reveal a potential GLP1R-mediated mechanism through which the alBST exerts inhibitory control over the endocrine HPA axis.

## Introduction

In rats and mice, physiological and behavioral stress responses are elicited and/or enhanced by increasing glucagon-like peptide-1 (GLP1) signaling within the brain, and are attenuated by reducing central GLP1 signaling (vanDijk and Thiele, 1999)(Kinzig et al., 2003)(Maniscalco et al., 2013)(Maniscalco et al., 2015)(Holt and Trapp, 2016)(Maniscalco and Rinaman, 2017)(Holt et al., 2019). GLP1 receptors (GLP1R) are expressed in multiple stress-related brain regions that receive direct axonal input from GLP1-positive neurons in the caudal nucleus of the solitary tract (NST) and adjacent intermediate reticular formation (Cork et al., 2015)(Heppner et al., 2015). These target regions include the anterior lateral bed nucleus of the stria terminalis (alBST), a limbic forebrain structure implicated in the central control of autonomic, neuroendocrine, and behavioral outflow (Dong and Swanson, 2006a)(Dong and Swanson, 2006b)(Crestani et al., 2013)(Johnson et al., 2016)(Johnson et al., 2019).

In recent years, rodent models have been used to examine how hindbrain GLP1 neural inputs to the alBST modulate the ability of acute stress to suppress food intake, promote anxiety-like behavior, and increase plasma levels of stress hormone [i.e., corticosterone (cort)] (Williams et al., 2018)(Zheng et al., 2019). We reported that virally-mediated suppression of GLP1R mRNA translation in the alBST has anxiolytic behavioral effects in rats, but paradoxically prolongs the plasma cort response to acute stress (Zheng et al., 2019). We interpreted the latter result as evidence that GLP1 signaling within the alBST normally engages an inhibitory “brake” on stress-induced activation of the hypothalamic-pituitary-adrenal (HPA) axis. This interpretation was bolstered by our finding that GLP1R-expressing neurons in the ventral alBST are GABAergic (Zheng et al., 2019), consistent with evidence that a GABAergic projection pathway from the ventral alBST to the paraventricular nucleus of the hypothalamus (PVN) serves to restrain stress-induced activation of the HPA axis (Radley et al., 2009)(Johnson et al., 2016). Further, GLP1R-expressing neurons in the dorsal and/or ventral alBST send axonal projections to the PVN and other stress-related brain regions in mice (Williams et al., 2018). In the same mouse study, whole-cell patch clamp recordings conducted in *ex vivo* slices revealed that bath-applied GLP1 evokes direct depolarizing or hyperpolarizing postsynaptic responses in GLP1R-expressing neurons in both dorsal and ventral portions of the alBST (Williams et al., 2018), evidence for complex GLP1R-mediated neural responses within this limbic forebrain structure. However, as neither the subnuclear distribution nor the projection targets of identified GLP1-responsive alBST neurons was examined, it remains unclear whether and how GLP1 affects the electrophysiological properties of PVN-projecting neurons within the ventral alBST.

Based on the evidence summarized above, we recently speculated that GLP1R is expressed by PVN-projecting GABAergic neurons within the ventral alBST, and hypothesized that GLP1 directly depolarizes/activates these inhibitory projection neurons (Zheng et al., 2019). The present study was designed to test this hypothesis directly in rats. In the first experiment, a fluorescent retrobead labeling strategy was combined with whole-cell patch recordings in *ex vivo* slices to identify PVN-projecting neurons within the ventral alBST, and to examine responses of these neurons to bath application of a specific GLP1R agonist. In the second experiment, a more sensitive retrograde tracer was combined with RNAscope fluorescent *in situ* hybridization to document the neurotransmitter phenotype of PVN-projecting neurons located in distinct subregions of the ventral alBST and adjacent parastrial nucleus (PS) of the hypothalamic preoptic region, and to determine whether these PVN-projecting neurons express mRNA for GLP1R. In a final follow-up experiment, retrograde labeling was combined with *in situ* hybridization to examine GLP1R mRNA expression in dorsal alBST neurons that provide axonal input to the ventral alBST.

## Methods

All animal procedures were conducted in accordance with the National Institutes of Health Guide for the Care and Use of Laboratory Animals (2011), and were approved by the Institutional Animal Care and Use Committee at the University of Pittsburgh and/or Florida State University.

### Electrophysiological analysis of PVN-projecting neurons in the ventral alBST

#### Retrograde labeling

PVN-projecting neurons were retrogradely labeled for subsequent collection of electrophysiological data in *ex vivo* slices. Young adult male Sprague-Dawley rats (Harlan; 180-190 g BW) were anesthetized by isoflurane inhalation (1–3% in oxygen; Halocarbon Laboratories) and fixed into a stereotaxic device in the flat skull position. Undiluted Red IX Retrobeads (LumaFluor, Inc. USA) were delivered by pressure (200nl/injection, 100 nl/min) into the PVN bilaterally (coordinates from bregma: −1.90 mm posterior, 0.30 mm lateral, −7.90 mm ventral) using a glass micropipette (~20 μm outer tip diameter) connected to a 10 μl Hamilton syringe controlled by a digital stereotaxic microinjector (catalog #QSI 53311, Stoelting). After each injection, the micropipette was left in place for 5 min to minimize tracer backflow. After the injector was removed, the skin over the skull was closed with 4-0 nylon sutures. Rats were injected subcutaneously with ketofen (2mg/kg BW) and buprenorphine (0.03 mg/kg BW) and were returned to their home cages after full recovery from anesthesia.

#### Whole-cell patch recordings

One week after stereotaxic Retrobead tracer injection into the PVN, rats were deeply anesthetized with isoflurane (5% in oxygen via inhalation) and then decapitated. The brain was quickly removed and immersed in ice-cold pre-oxygenated artificial cerebrospinal fluid (ACSF). A tissue block containing the anterior BST was excised, and coronal slices (350-400 μm thick) were cut with a vibratome (Leica VT1000S, Leica, Germany). Visual landmarks including the anterior commissure, lateral ventricles, and optic chiasm were used to select slices through the appropriate rostrocaudal level of the anterior BST [i.e., approximately 0.2-0.5 mm caudal to bregma, based on the most recent version of Swanson’s rat brain atlas (Swanson, 2018); see **Figure 1**]. Slices were incubated at 37°C for 0.5-1 h, then kept at room temperature before being transferred to a recording chamber perfused with oxygenated ACSF (95% O_2_/5% CO_2_; pH 7.25-7.3) at 31-32°C. ACSF solution composition (in mM) was 126 NaCl, 2.5 KCl, 1.25 NaH_2_PO_4_, 1 MgSO_4_, 2 CaCl_2_, 24 NaHCO_3_, and 10-20 glucose.

**Figure 1.**
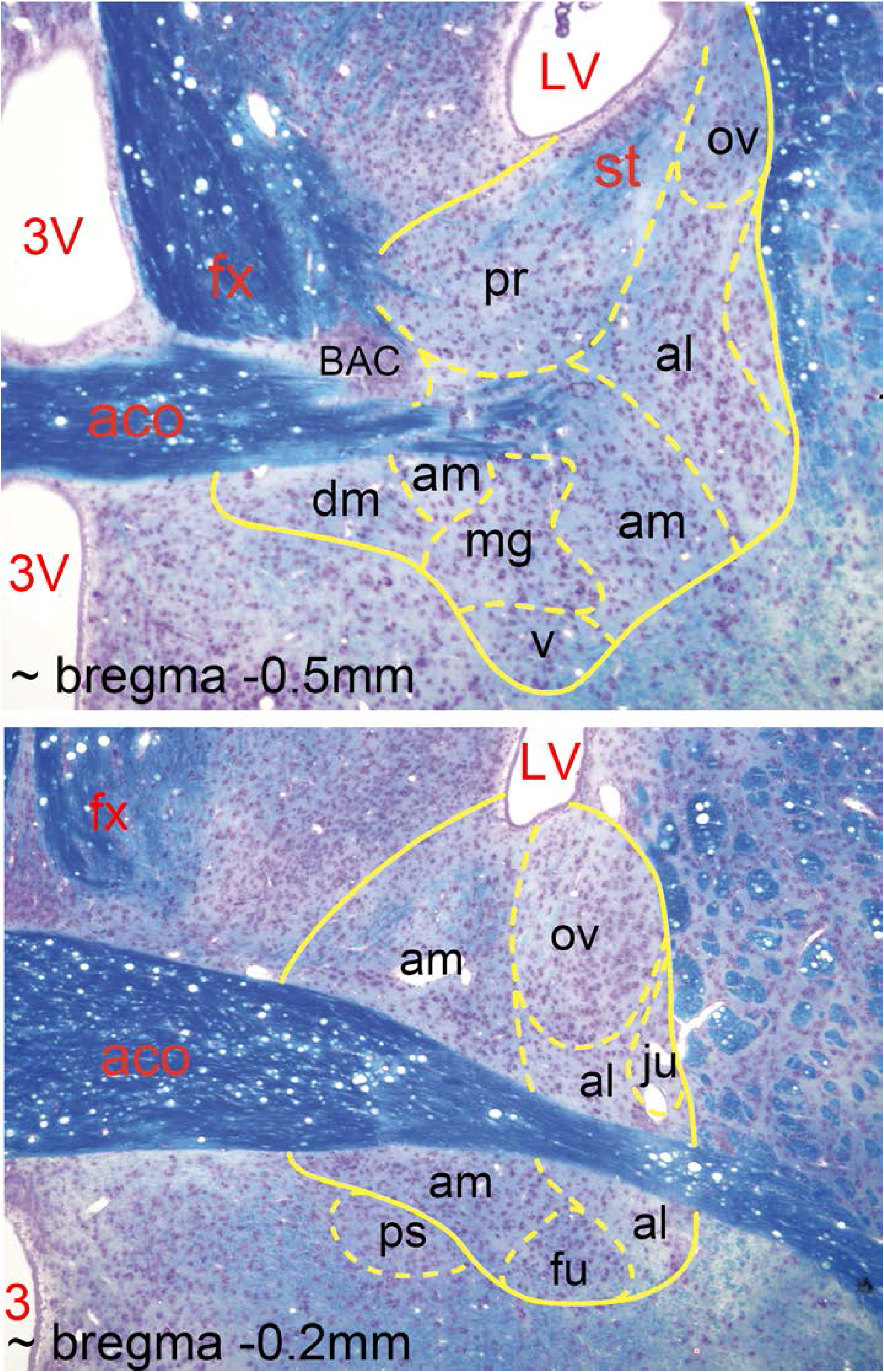
Subnuclear organization of the rat alBST. Klüver-Barrera-stained coronal tissue sections through two representative rostrocaudal levels of the alBST. Myelinated fibers are blue, neuron cell bodies are pink/purple. Images are adapted from our previous publication (Bienkowski et al., 2013), with BST subnuclei outlined in yellow and labeled according to the most recent version of Swanson’s rat brain atlas (Swanson, 2018). 3V, third ventricle; aco, anterior commissure; al, anterolateral subnucleus; am, anteromedial subnucleus; BAC, bed nucleus anterior commissure; dm, dorsomedial subnucleus, fu, fusiform subnucleus; fx, fornix; ju, juxtacapsular subnucleus; LV, lateral ventricle; mg, magnocellular subnucleus; ov, oval subnucleus; pr, principal subnucleus; ps, parastrial subnucleus of the hypothalamus; st, stria terminalis; v, ventral subnucleus. The approximate (~) rostro-caudal level of each section (relevant to bregma) is indicated; the more caudal section is at the top, and the more rostral section at the bottom.

A Zeiss Axioskop microscope (Carl Zeiss, Inc., Thornwood, NY) equipped with a 40x water immersion objective and a digital video camera (CoolSnap, Photometrics, Tucson, Az) was used to visualize retrogradely-labeled neurons for whole-cell patch recordings. Within each slice, the ventral region of the alBST was identified by its medial-lateral position (i.e., vertically aligned with the lateral ventricle, medial to the internal capsule), by its close ventral proximity to the anterior commissure, and by the presence of red retrobead-labeled neurons. Patch electrodes were filled with an internal solution containing (in mM): 105 Cs-gluconate, 2 MgCl_2_, 10 NaCl, 10 HEPES, 10 phosphocreatine, 4 ATP-Mg, 0.3 GTP, and 10 BAPTA; pH 7.25. In addition, Alexa 568 (0.075%; Molecular Probes, Eugene, OR) was added to the patch solution to fill each recorded neuron for later morphological identification, as previously described (Povysheva et al., 2006). Electrodes had 5-10 MΩ open-tip resistance. Voltage and current recordings were performed with a Multi-Clamp 700A amplifier (Axon Instruments, Union City, CA). Voltage recordings were performed in bridge-balance mode. Signals were filtered at 2 KHz, and acquired at a sampling rate of 10 kHz using a Digidata 1440 digitizer and Clampex 10.2 software (Molecular Devices Corporation, Sunnyvale, CA). Access resistance and capacitance were compensated on-line. Access resistance typically was 10-20 MΩ and remained relatively stable during experiments (≤ 30% increase) for cells included in data analysis. Membrane potential was corrected for the liquid junction potential of −13 mV.

To characterize neuronal membrane properties, hyper- and depolarizing current steps were applied for 500 ms in increments of 5 to 10 pA at 0.5 Hz. Input resistance was measured from the slope of a linear regression fit to the voltage-current relation in a voltage range hyperpolarized from resting potential. The membrane time constant was determined by single-exponential fitting to the average voltage responses activated by hyperpolarizing current steps of 5-15 pA. Action potential (AP) properties were quantified using the first action potential evoked through application of depolarizing current steps. AP threshold was measured at the level of voltage deflection exceeding 10 mV/1 ms. Peak AP amplitudes of the AP and afterhyperpolarization were measured relative to AP threshold. AP duration was measured at the base (AP threshold level). Action potential frequency was calculated in Hz as a ratio between number of action potentials and current step duration at 60 pA above the rheobase.

The synthetic GLP1 analogue Exendin-4 (Ex-4; 200-600 nM; Bachem, Torrance, CA) was bath-applied to activate GLP1Rs, while the specific antagonist Exendin-9 (Ex-9; 900 nM; Bachem) was used to block GLP1Rs (Goke et al., 1993)(Thorens et al., 1993). Additional pharmacological agents were bath applied at the following concentrations: tetrodotoxin (TTX; 0.5 μM; Sigma) to block voltage-gated Na^+^ channels; AP-5 (D-2-amino-5-phospho-pentanoic acid; 50 μM; Ascent Scientific LTD, Bristol, UK) to block NMDA (N-methyl-D-aspartate) receptors; NBQX (2,3-dihydroxy-6-nitro-7-sulfamoylbenzo(F)quinoxaline; 20 μM; Ascent Scientific) to block AMPA (α-amino-3-hydroxy-5-methyl-4-isoxazolepropionic acid) and kainate receptors; gabazine (10 μM; Ascent Scientific LTD, Bristol, UK) to block GABA-A receptors.

Patch recordings in identified ventral alBST neurons were made in the presence or absence of Ex-4. Spontaneous inhibitory post-synaptic currents (sIPSCs) were recorded at a holding potential of +12 mV in the presence of NBQX and AP-5. Miniature inhibitory post-synaptic currents (mIPSCs) were recorded at a holding potential of +12 mV in the presence of TTX to inhibit action potential-mediated inhibitory postsynaptic currents. Spontaneous excitatory post-synaptic currents (sEPSCs) were recorded at a holding potential of −70 mV. Miniature excitatory post-synaptic currents (mEPSCs) were recorded at a holding potential of −70 mV in the presence of TTX to inhibit action potential-mediated excitatory postsynaptic currents.

Spontaneous and miniature events were analyzed using the MiniAnalysis Program (Synaptosoft, Decatur, GA). Peak events were first detected automatically using an amplitude threshold of 1.5 times the average RMS noise, which approximated 3 pA for recordings at a holding potential of −70mV. More than 500 events per cell were included in each analysis.

#### Morphological data analysis

Neurons were filled with Alexa 568 during whole-cell recordings. A subset of these were maintained for at least 30 min to ensure extensive dendritic labeling. After recording, slices were fixed in ice-cold 4% paraformaldehyde for at least 72 h, then transferred into an anti-freeze solution (ethylene glycol and glycerol in 0.1 M phosphate buffer) and stored at −20C. A few representative labeled neurons were reconstructed three-dimensionally using an Olympus Fluoview BX61 confocal microscope and Fluoview software (Olympus America Inc, Melville, NY).

#### Statistical analysis of electrophysiological data

Unless otherwise specified, one-way ANOVA and post-hoc two-tailed paired t-tests were used for between-group comparisons. Values are presented as mean ± SEM. Statistical tests were performed using Excel (Microsoft Corp., Redmond, WA). Effects were considered significant when p<0.05.

### Molecular phenotyping of projection-specific alBST neurons

#### Retrograde labeling

In a separate experiment, PVN-projecting neurons were retrogradely labeled and visualized using immunocytochemical detection of cholera toxin beta (CTB) neural tracer combined with dual *in situ* hybridization to analyze cellular expression of GLP1R plus Vgat mRNAs to identify GABAergic neurons, or GLP1R plus Vglut2 mRNAs to identify glutamatergic neurons. For this, adult male Sprague-Dawley rats (n=3; 225-250 g BW) were anesthetized by isoflurane inhalation (1-3% in oxygen; Halocarbon Laboratories) and placed into a stereotaxic device in the flat skull position. A pulled glass pipette (~20 μm outer tip diameter) was attached to the arm of the stereotaxic apparatus. A solution of 0.25% CTB (List Biological Labs, Campbell, CA, USA) in sodium phosphate buffer (pH 7.5) was backfilled through the pipette tip using negative pressure, then a wire connected to a current source (Stoelting) was inserted into the tracer solution. During descent of the glass pipette into the brain, a −0.5 μA retaining current was used to minimize molecular diffusion of tracer from the pipette tip. CTB was iontophoresed unilaterally into the PVN (from bregma: 1.90 mm posterior, 0.30 mm lateral, −7.90 mm ventral) using a 7 s on/off pulsed current of +5 μA for 10 min. Five minutes after the end of iontophoresis, a −0.5 μA retaining current was held as the pipette was withdrawn, and the skin over the skull was closed with 4-0 nylon sutures. Rats were injected subcutaneously with ketofen (2mg/kg BW) and buprenorphine (0.03 mg/kg BW) and were returned to their home cages after recovery from anesthesia.

In a separate follow-up experiment, CTB was iontophoresed unilaterally as described above, but with the tracer delivery site targeting the ventral alBST (instead of PVN) in one adult male rat (from bregma: 0.3 mm posterior, 1.7 mm lateral, and 7.4 mm ventral). The goal was to determine whether dorsal alBST neurons known to innervate the ventral alBST (Bienkowski and Rinaman, 2013) express GLP1R, and whether these neurons have a GABAergic or glutamatergic phenotype.

Ten to twelve days after CTB delivery into PVN or alBST, rats were anesthetized with a lethal dose of sodium pentobarbital (Fatal Plus, 100 mg/kg; Butler Schein, Columbus, OH) and then transcardially perfused with physiological saline followed by a fixative solution containing 4% paraformaldehyde, 1.4% lysine and 0.3% sodium metaperiodate in 0.1 M sodium phosphate buffer (McLean and Nakane, 1974). Brains were extracted from the skull, blocked and postfixed overnight at 4°C, and cryoprotected in 20% sucrose solution for 24-72 hrs. Brains were then sectioned coronally (35 μm) using a freezing sliding microtome. Sections were collected sequentially into six adjacent sets and stored in cryopreservant solution (Watson et al., 1986) at −20°C for subsequent processing.

#### Dual immunocytochemistry and in situ hybridization

Immunofluorescence was combined with RNAscope *in situ* hybridization in order to assess GLP1R mRNA expression together with expression of either Vgat or Vglut2 mRNA in retrogradely labeled (i.e., CTB-immunopositive) projection neurons.

##### RNAscope in situ hybridization

mRNAs were identified using Rn-Glp1r probe (315221, Accession No. NM_012728.1, Target Region: 292-1166), Vgat probe (Rn-Slc32a1) and Vglut2 probe (Rn-Slc17a6), using RNAscope^®^ Multiplex Fluorescent Reagent Kit v2 (323100). Selected tissue sections containing the anterior BST were removed from cryopreservant solution and washed for 1 □hr in four changes of 0.1M PB. Sections were pretreated with H_2_O_2_ solution (ACD 322335) for 30 min followed by 4 × 8 min rinses in 0.1M PB. Pretreated sections were then mounted from 0.01M Tris buffer (TB) onto Gold Seal™ UltraStick™ Adhesion Microscope Slides (3039-02; ThermoFisher Scientific, Waltham, MA, USA) and air-dried at room temperature for 1 □hr. Slides were dipped into 100% ethanol for 10 sec and then air-dried for 2-3 hours before creating a hydrophobic barrier around each tissue section using a Pap Pen (195506; ThermoFisher Scientific). The barrier was allowed to completely dry at room temperature overnight before proceeding to the next step. Unless otherwise noted, incubations were performed at 40°C within a HybEZTM Oven, using the HybEZTM Humidity Control Tray. For each section, two to four drops of each reagent solution were used to cover tissue sections, followed by three 3-min rinses in 1× washing buffer at room temperature. Sections were incubated with Protease IV (322336) for 20-25 min at room temperature, then washed in 0.01M TB four times (1 min/wash). Sections were then incubated in a cocktail of two probes (Rn-Glp1r plus either Rn-Vgat or Rn-Vglut2 for 2 hr), followed sequentially by amplification steps with Amp1 (323101) for 30 min, Amp2 (323102) for 30 min, and Apm3 (323103) for 15 min. GLP1R mRNA and Vgat mRNA or Vglut2 mRNA were then labeled with fluorophore-conjugated Tyramine plus (TSAP) sequentially according to the manufacturer’s protocol. For GLP1R mRNA, sections were incubated in channel 1-specific HRP (323104) for 15min and Cy3-TSAP (1.5K, tyramine signal amplification plus, NEL744E001KT; PerkinElmer Waltham, MA, USA) for 30 min followed by a 15 min incubation in HRP blocker (323107). Sections were then incubated in channel 2-specific HRP (323105) for Vgat mRNA or Vglut2 mRNA for 15 min followed by Cy5-TSAP (1.5K) for 30 min. After a final washing buffer rinse, slides were rinsed in 0.1M PB for 15 min before proceeding to immunofluorescent labeling of CTB.

##### Immunofluorescent labeling

:Incubations were performed at room temperature using the HybEZTM Humidity Control Tray with gentle agitation in a shaker. Sections already processed for RNAscope labeling were incubated in goat anti-CTB antiserum (1:5K, List Biological Labs, cat #703) for 24 hours, rinsed for 40 min in 4 changes of 0.1M PB, and then incubated in Alexa 488-conjugated donkey anti-goat secondary antibody (1:400, Jackson ImmunoResearch) for 3 hours. After a final rinse in 0.1M PB, sections were air-dried overnight, dehydrated/defatted in graded ethanols and xylene (2 min each), and coverslipped using Cytoseal 60 (VWR).

##### Image acquisition

Low magnification images were collected using a bright-field and epifluorescent Keyence microscope (catalog #BZ-X700). Higher-resolution images were acquired using a Leica TCS SP8 Confocal Microscope with a 20X air-objective and a 100X oil-objective. Alexa 488 and Cy3 were excited using 488 nm and 552 nm OPSL lasers, respectively, and Cy5 using a 638 nm Diode laser. Confocal images were obtained using Leica LAS version 4.0 image software, with images of each tissue region collected sequentially for each fluorophore to avoid signal contamination. Leica imaging software was used to generate Z plane projections and 3D rotatable maximum intensity projections.

## Results

### Electrophysiological analysis of PVN-projecting neurons in the ventral alBST/PS

Electrophysiological data were collected from PVN-projecting (i.e., tracer-labeled) neurons within the ventral alBST. Given the proximity of the hypothalamic PS to the ventral alBST at rostrocaudal levels used to prepare slices for electrophysiology (see **Fig. 1**), and given that ventral alBST and PS neurons similarly project to the PVN in rats (Simerly and Swanson, 1988)(Thompson, 2003)(Dong and Swanson, 2006b), we assume that tracer-labeled neurons selected for patch recording included both PS and alBST neurons. Additional recordings were made in non-tracer-labeled alBST/PS neurons located closely adjacent to tracer-labeled neurons. **Table 1** summarizes the membrane properties of tracer-labeled and non-labeled neurons recorded before bath application of the specific GLP1R agonist Ex-4 in concentrations ranging from 200-600 nM. Despite significant main effects of Ex-4 on the membrane properties of tracer-labeled, PVN-projecting neurons (described below), these effects were independent of Ex-4 concentration. For this reason, data collected from multiple slices and neurons within each experiment are combined in Figures 1–4, regardless of Ex-4 concentration. In each figure, the Ex-4 concentration used during data collection in individual representative neurons is indicated.

**Table 1.** Membrane properties of tracer-labeled and non-labeled alBST/PS neurons. *Abbreviations:* AHP, afterhyperpolarization; AP, action potential; APT, action potential threshold; RMP, resting membrane potential

| **Membrane property** | **Tracer-labeled**<br>**(*n*=11 neurons**<br>**from 7 rats)** | **Non-labeled**<br>**(*n*=10 neurons**<br>**from 5 rats)** |
| --- | --- | --- |
| RMP, mV | -58.3±4.3 | -57.4±3.7 |
| APT, mV | -36.8±5.9 | -37.8±3.9 |
| AHP amplitude, mV | -13.6±5.2 | -10.5±3.9 |
| Spike frequency, Hz | 18.3±9.1 | 21.2±14.0 |
| AP amplitude, mV | 59.7±2.9 | 57.4±7.6 |
| AP duration, ms | 2.4±0.8 | 2.0±0.6 |
| Input resistance, MΩ | 555.8±177.8 | 421.6±205.7 |
| Time constant, ms | 30.7±12.3*<br>* *p* < 0.05 vs.<br>*non-labeled* | 22.1±5.1 |

#### Ex-4 depolarizes the baseline membrane potential of PVN-projecting neurons in the ventral alBST/PS

PVN-projecting neurons in each slice were identified by the presence of red retrobead fluorescent labeling (see **Fig. 2E**). A subset of labeled neurons in the ventral alBST/PS were selected for whole-cell patch recording (n=11 neurons from 7 rats). The membrane properties of bead-labeled neurons before bath application of Ex-4 are reported in **Table 1**. Only 2 out of 11 neurons displayed spontaneous action potentials.

**Figure 2.**
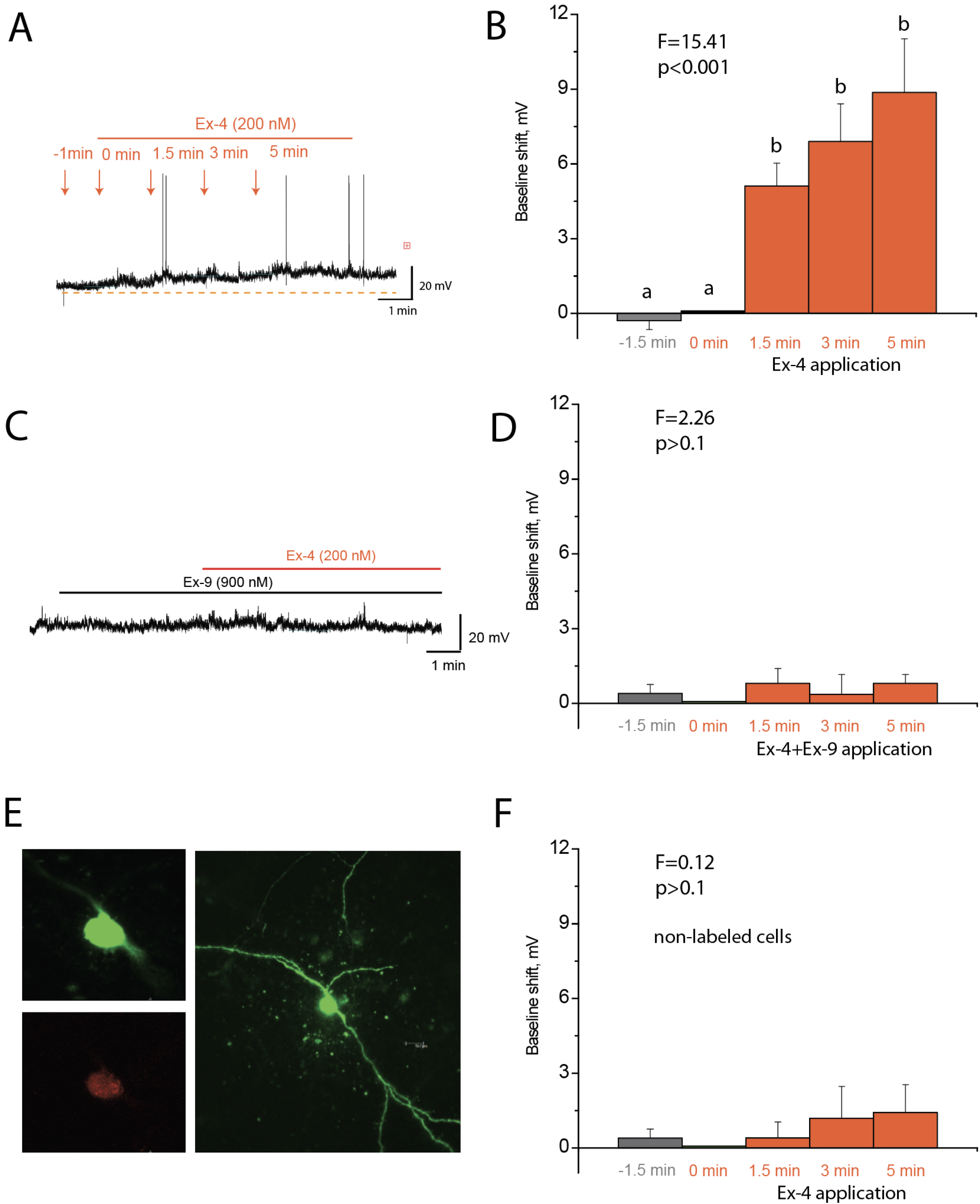
Shift in baseline membrane potential produced by Ex-4. **(A)** Baseline recording in a representative PVN-projecting neuron within the anterior vlBST/PS before and after bath application of the specific GLP1R agonist, Ex-4 (200 nM). **(B)** Summary data indicating that Ex-4 (200-600 nM) produced a depolarizing shift of the baseline membrane potential in PVN-projecting cells (n=11 neurons from 7 rats), which was significant from 1.5 to 5 min after bath application. **(C)** Baseline recording in a representative PVN-projecting neuron before and after bath application of Ex-4 (200 nM) in the presence of 900 nM Ex-9, a specific GLP1R antagonist. **(D)** Summary data indicating that Ex-4 (200-600 nM) effects on baseline membrane potential in PVN-projecting neurons (n=6 cells from 4 rats) are occluded by Ex-9. **(E)** A representative PVN-projecting neuron double-labeled with retrobeads (red, bottom left) and Alexa 488 dye (green) applied during neural recording. The same neuron is confocally reconstructed (right panel) to display its morphology. **(F)** Ex-4 (200-600 nM) had no effect on baseline membrane potential in non-labelled cells (n=10 neurons from 5 rats) located in the vicinity of PVN-projecting neurons.

We next examined the effect of bath-applied Ex-4 on labeled neurons. One-way ANOVA revealed that Ex-4 produced a substantial upward shift in baseline membrane potential during a 5 min application period, which induced spiking in 3/11 neurons. Compared to baseline membrane potential recorded 1.5 min before and at the time of Ex-4 application, the upward shift was significant at 1.5, 3, and 5 minutes post-application (**Fig. 2A, 2B**). While recording from labeled neurons contained in other slices (n=6 neurons from 4 rats), we examined whether the depolarizing effect of Ex-4 is mediated through activation of GLP1Rs. For this, the GLP1R antagonist Ex-9 was bath-applied a few minutes before application of Ex-4. While Ex-9 by itself had no effect on baseline membrane potential, it completely occluded the effect of subsequently administered Ex-4 (**Fig. 2C, 2D**).

In several slices, whole-cell recordings were made in additional non-labeled neurons that were positioned in close proximity to labeled neurons in the ventral alBST/PS (n=10 neurons from 5 rats). These nearby non-tracer-labeled neurons possessed intrinsic membrane properties similar to those displayed by labelled neurons; the only documented exception was a significantly slower time constant displayed by tracer-labeled vs. non-labeled neurons (**Table 1**). Ex-4 failed to generate detectable shifts in baseline membrane potential of non-labeled neurons during the 5-minute application period (**Fig. 2F**).

#### Synaptic blockers eliminate Ex-4 depolarizing effects on membrane potential

To determine whether Ex-4-induced membrane depolarization is due to effects on the intrinsic membrane properties of PVN-projecting neurons within the ventral alBST/PS (n=10 neurons from 5 rats), Ex-4 was applied a few minutes after bath application of synaptic blockers. For this, AMPA and kainate receptors were blocked with NBQX, NMDA receptors were blocked with AP-5, and GABA-A receptors were blocked with gabazine. Under these conditions of synaptic blockade, Ex-4 did not significantly alter baseline membrane potential in recorded neurons (**Fig. 3A, 3B**). Thus, the depolarizing effect of Ex-4 on PVN-projecting neurons (**Fig. 2**) depended on modulation of their presynaptic inputs, rather than directly affecting intrinsic ion channel conductances.

**Figure 3.**
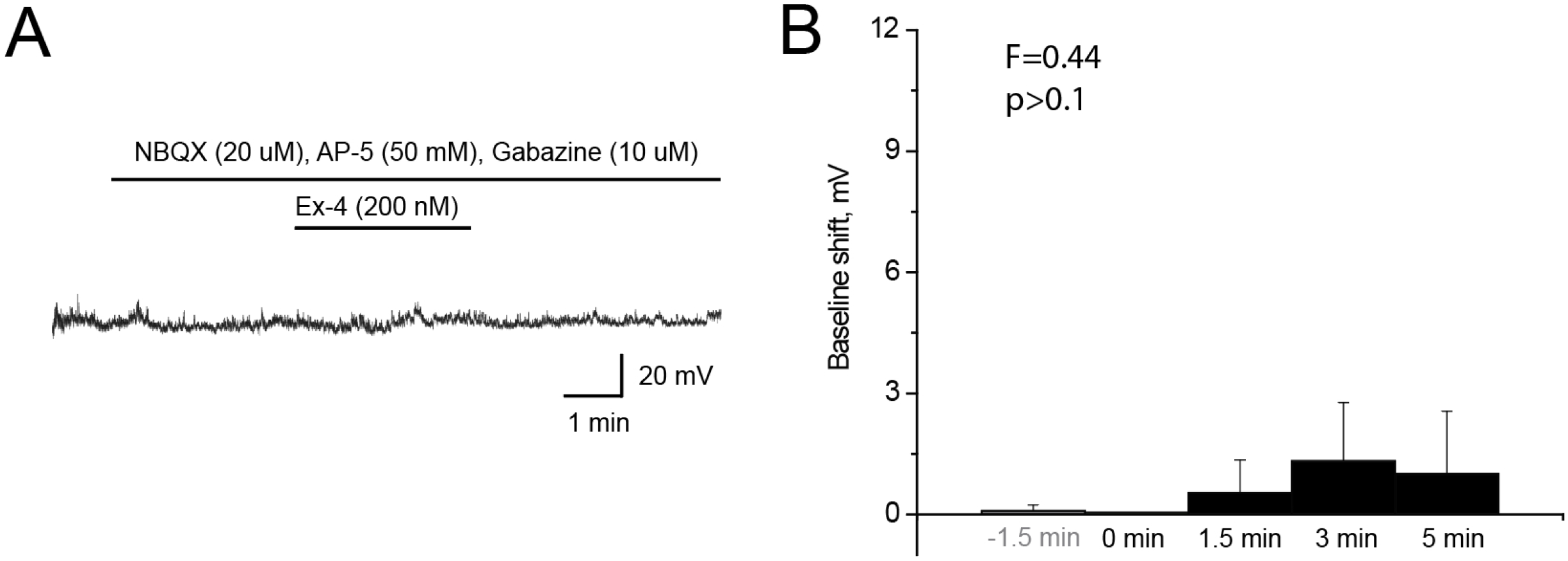
Synaptic blockers eliminate Ex-4 effects on baseline membrane potential in PVN-projecting neurons. **(A)** Baseline recording in a representative PVN-projecting neuron in the anterior vlBST/PS, recorded before and after bath application of Ex-4 (200 nM) in the presence of synaptic blockers. **(B)** Summary data indicating that Ex-4 (200-600 nM) effects on baseline membrane potential (n=10 neurons from 5 rats) were eliminated by prior bath application of synaptic blockers.

#### Ex-4 reduces inhibitory inputs to PVN-projecting neurons in the ventral alBST/PS

To assess Ex-4 effects on inhibitory responses in PVN-projecting neurons, sIPSCs were recorded in voltage-clamp mode at a holding potential of +12 mV in the presence of NBQX and AP-5 to block glutamatergic receptor signaling. Subsequent bath application of Ex-4 significantly reduced both the frequency and amplitude of sIPSCs recorded in PVN-projecting neurons (n=7 neurons from 4 rats; **Fig. 4A, 4B, 4C**). These sIPSCs could result from action potential-dependent network activity within the slice, and/or action potential-independent transmitter release. To evaluate these possibilities, TTX was applied to block voltage-gated Na+ channels (and hence, neural firing) within the slice, leaving only action potential-independent mIPSCs. The frequency of mIPSCs represents presynaptic GABA release, whereas mIPSC amplitude is a property of postsynaptic receptors. When PVN-projecting cells were recorded in the presence of TTX (n=13 neurons from 6 rats), Ex-4 reduced the frequency of mIPSCs (**Fig. 4D**) but did not change their amplitude (**Fig. 4E**), supporting a presynaptic effect of Ex-4 to reduce inhibitory GABA input to PVN-projecting neurons.

**Figure 4.**
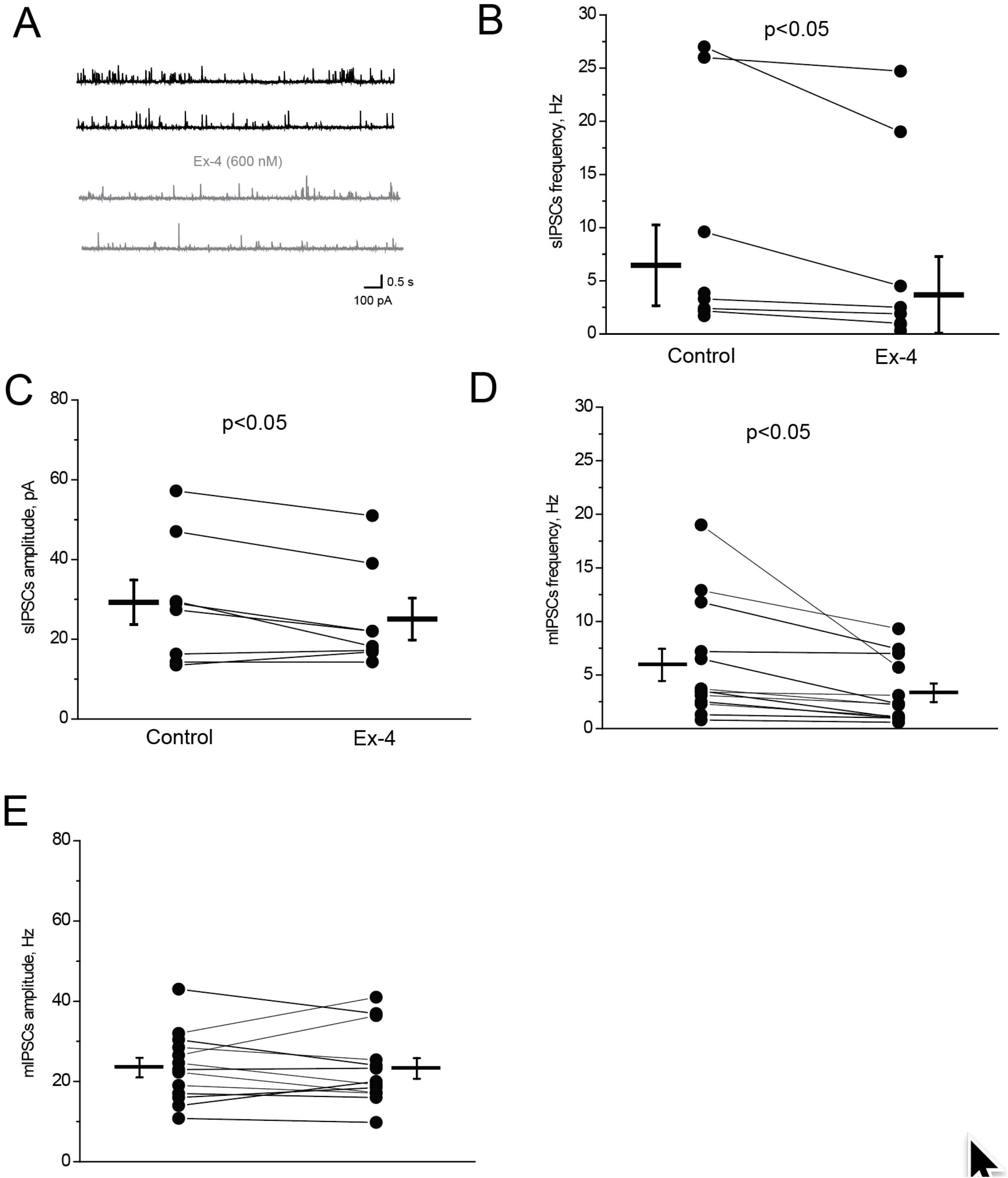
Ex-4 reduces inhibitory inputs to PVN-projecting neurons. **(A)** sIPSCs recorded in a representative PVN-projecting neuron in the anterior vlBST/PS before (top traces) and after (bottom traces) bath application of 600 nM Ex-4. **(B)** Summary data indicating that Ex-4 decreased sIPSC frequency in PVN-projecting neurons (n=7 neurons from 4 rats). **(C)** Summary data indicating that Ex-4 decreased sIPSC amplitude in the same PVN-projecting neurons. **(D)** In the presence of TTX, Ex-4 decreased mIPSC frequency but not mIPSC amplitude **(E**) in PVN-projecting neurons (n=13 neurons from 6 rats).

#### Ex-4 increases excitatory inputs to PVN-projecting neurons in the ventral alBST/PS

To assess Ex-4 effects on sEPSCs, PVN-projecting neurons were recorded in voltage-clamp mode at a holding potential of −70 mV, a condition under which sIPSCs are undetectable (given the −67 mV reversal potential for GABA_A_ responses). When cells were recorded under these conditions (n=8 neurons from 4 rats), bath-applied Ex-4 increased the frequency but not the amplitude of sEPSCs (**Fig. 5A, 5B, 5C**), supporting a presynaptic effect of Ex-4 to increase excitatory inputs to the recorded neurons. To confirm this, TTX was bath-applied to isolate action potential-independent mEPSCs. In PVN-projecting neurons recorded under these conditions (n=6 cells from 3 rats), Ex-4 altered neither mEPSC frequency (**Fig. 5D**) nor amplitude (**Fig. 5E**).

**Figure 5.**
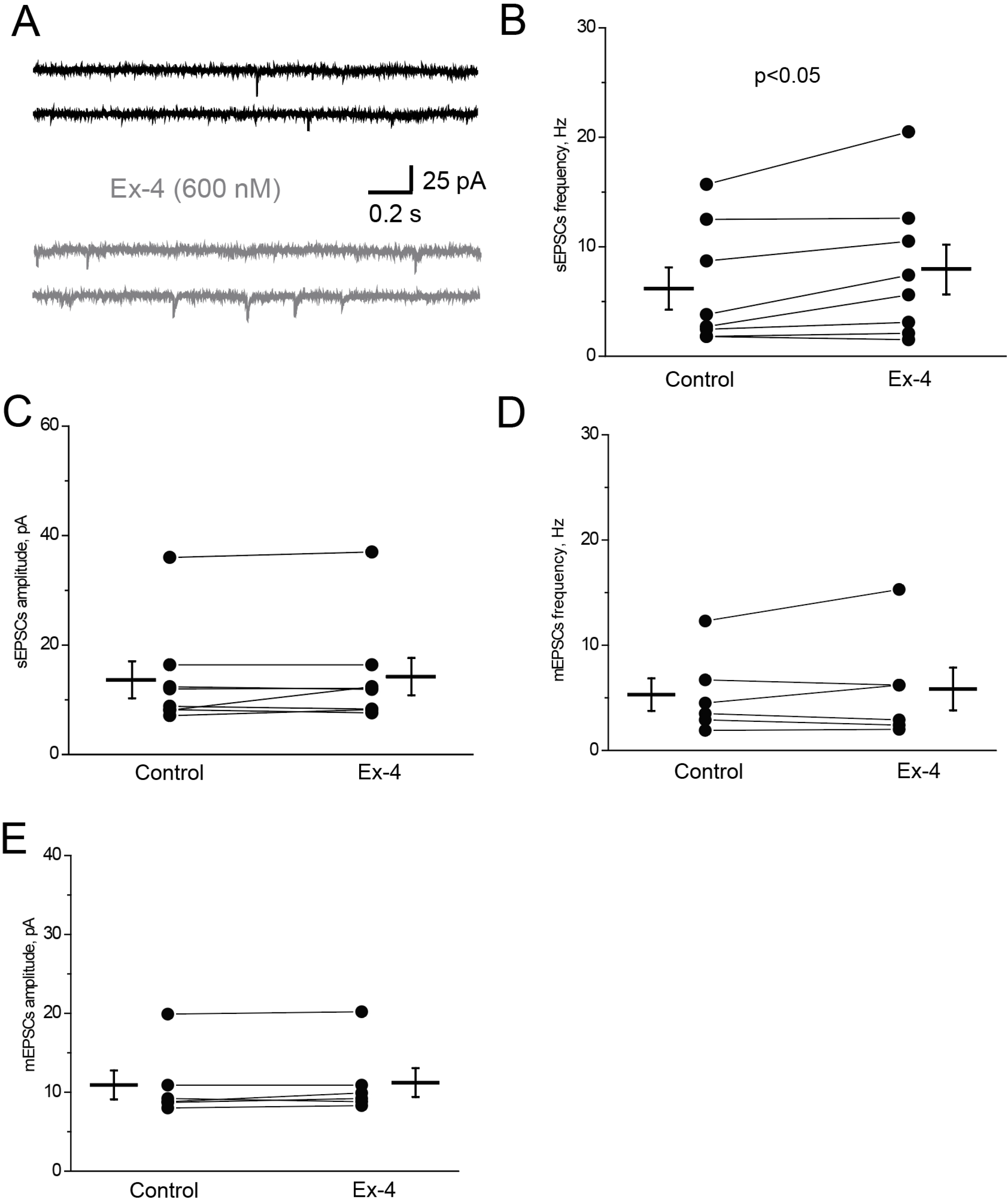
Ex-4 increases excitatory inputs to PVN-projecting neurons. **(A)** sEPSCs recorded in a representative PVN-projecting neuron within the anterior vlBST/PS before (upper trace) and after (lower trace) bath application of 600 nM Ex-4. **(B)** Summary data demonstrate that Ex-4 increased sEPSC frequency but not sEPSC amplitude **(C)** in PVN-projecting neurons (n=8 neurons from 4 rats). **(D)** In the presence of TTX, Ex-4 had no significant effect on either mEPSC frequency or mEPSC amplitude **(E).**

#### Molecular phenotyping of projection-specific neurons in the ventral alBST/PS

The slice electrophysiology results described above (**Figs. 2–5**) indicate that bath-applied GLP1R agonist (i.e., Ex-4) reduced inhibitory inputs and increased excitatory inputs to retrobead-labeled, PVN-projecting neurons within the ventral alBST/PS. Notably, labeled projection neurons recorded in the presence of TTX did not respond directly to Ex-4, suggesting that the majority (and perhaps all) PVN-projecting neurons lack functional GLP1Rs. These results were surprising, because we originally hypothesized that GLP1 would directly activate PVN-projecting neurons in the ventral alBST. To confirm the electrophysiological findings, a separate experiment using a different neural tracer (i.e., CTB) and immunocytochemical amplification of retrograde labeling was conducted to more fully and accurately localize PVN-projecting neurons within the ventral alBST and hypothalamic PS. Next, to examine local sources of afferent input to the ventral alBST and to determine whether they express GLP1R mRNA, a final follow-up experiment used CTB tracing from the ventral alBST to identify retrogradely-labeled neurons within the dorsal alBST at the same rostrocaudal level (i.e., likely contained within *ex vivo* slices used for electrophysiological analyses). In each CTB neural tracing experiment, tracer immunolabeling was combined with RNAscope-based *in situ* hybridization to colocalize neural expression of mRNA for GLP1R along with Vgat or Vglut2.

#### Distribution of PVN-projecting neurons

Following iontophoretic CTB tracer delivery into the hypothalamic PVN, retrogradely-labeled neurons were observed within many distinct regions of the forebrain and brainstem, consistent with results from previous studies reporting many sources of afferent input to the rat PVN (Sawchenko and Swanson, 1983)(Swanson and Sawchenko, 1980). The present analysis was limited to retrograde labeling in coronal tissue sections through the rostrocaudal level of the anterior BST (i.e., approximately 0.2-0.5 mm caudal to bregma; see **Fig. 1**), the level at which coronal *ex vivo* slices were taken for electrophysiological recordings in the first experiment. Consistent with previous reports (Dong et al., 2001)(Dong and Swanson, 2006b)(Lebow and Chen, 2016), CTB-positive neurons occupied ventral alBST regions corresponding to the location of the anteromedial, ventral dorsomedial, fusiform, and anterolateral subnuclei (**Figs. 6, 7**). At the more rostral levels included in analysis, an additional prominent cluster of retrogradely-labeled neurons was positioned slightly medial to the ventral alBST, corresponding to the location of the hypothalamic PS (see **Fig. 1, lower panel**). Unlike the BST, the PS does not receive input from the amygdala, and is thus considered to be part of the hypothalamic preoptic region rather than the BST. However, the PS and ventral alBST share many inputs and outputs, including a prominent projection to the PVN (Simerly and Swanson, 1988)(Thompson, 2003)(Dong and Swanson, 2006b). CTB-positive neurons also were present within the substantia innominata, and throughout the median preoptic and anteroventral periventricular nuclei of the anterior hypothalamus. It should be noted that tracer-labeled neurons in these locations were not included in *ex vivo* slice electrophysiology, because their positions did not meet the selection criteria (i.e., not close to the anterior commissure, not vertically aligned with the lateral ventricle).

**Figure 6.**
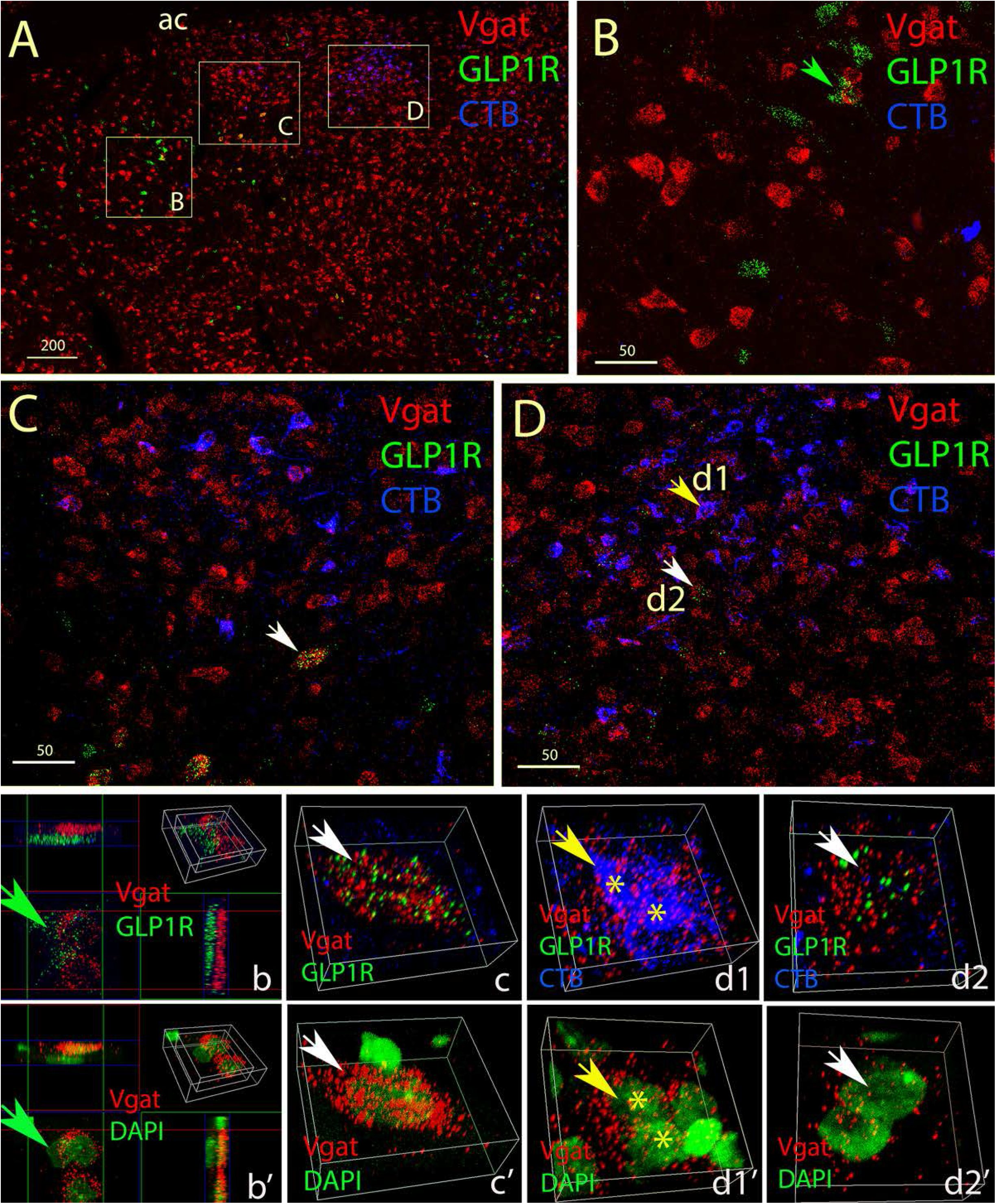
PVN-projecting neurons in the hypothalamic PS and in the ventral anterolateral and fusiform subnuclei of the alBST are predominantly GABAergic and do not express GLP1R. **(A)** Low magnification Z-max projection image of a coronal section through the ventral alBST (left hemisphere; approximately 0.2 mm caudal to bregma), showing the distribution of PVN-projecting CTB-positive (blue) neurons, neurons expressing Vgat mRNA (red), and neurons expressing GLP1R mRNA (green). Dorsal is towards the top, medial is towards the right. From lateral to medial, the boxed regions indicated by the labels B, C, and D are shown at higher magnification in the correspondingly labeled panels. ac, anterior commissure. Scale bar, 200 microns. **(B)** High magnification Z-max projection image from the most lateral boxed sub-region labeled B in panel A. This region corresponds to the ventrolateral edge of the anterolateral subnucleus of the BST, and also includes a portion of the adjacent substantia innominata. Many neurons in this field of view express GLP1R mRNA (green) or Vgat mRNA (red), but these two mRNA tags are not colocalized. A few CTB-positive neurons (blue) in the substantia innominata express neither Vgat nor GLP1R mRNA. The green arrowhead points out two adjacent Vgat+ neurons depicted at higher magnification in panels b, b’, with 3D-MIP (maximum intensity projections). A GLP1R+ neuron (green) is located adjacent to the two Vgat-expressing neurons (red). Panel b’ shows the same Vgat+ cells (red) imaged along with DAPI nuclear label (green). **(C)** High magnification Z-max projection image from the boxed sub-region labeled C in panel A, corresponding to a region containing the fusiform subnucleus of the BST. CTB labeling (blue) is imaged along with Vgat mRNA (red) and GLP1R mRNA (green). Many CTB-positive neurons are visible, all of which express Vgat mRNA. However, no CTB-labeled cells in this region express GLP1R mRNA. Several GLP1R+ neurons are present within the lower (more ventral) region in this field of view; most of these express Vgat mRNA. The white arrowhead in panel C points out a cell shown at higher magnification in panels c, c’, with 3D-MIP to demonstrate colocalization of Vgat and GLP1R mRNAs (c) and Vgat mRNA/DAPI (c’). **(D)** High magnification Z-max projection image from the most medial boxed sub-region labeled D in panel A, corresponding to the location of the hypothalamic PS. CTB labeling (blue) is imaged along with Vgat mRNA (red) and GLP1R mRNA (green). At this rostrocaudal level of the PS, most CTB-labeled neurons express Vgat mRNA, but none were observed to express GLP1R mRNA. Most of the GLP1R+ neurons visible within this region were observed to express Vgat mRNA. The yellow arrowhead (labeled d1) points out two CTB-positive cells shown at higher magnification in panel d1 (asterisks) with 3D-MIP to demonstrate their colocalization of Vgat mRNA (red). Panel d1’ shows the same field of view with Vgat mRNA labeling (red) imaged along with nuclear DAPI (green). The white arrowhead in panel D (labeled d2) indicates two non-tracer-labeled neurons, shown at higher magnification (3D-MIP) in panel d2 to demonstrate cellular colocalization of Vgat (red) and GLP1R mRNA (green). Intracellular colocalization of both mRNA labels is demonstrated in panel d2’, in which Vgat mRNA (red) is imaged along with nuclear DAPI (green).

**Figure 7.**
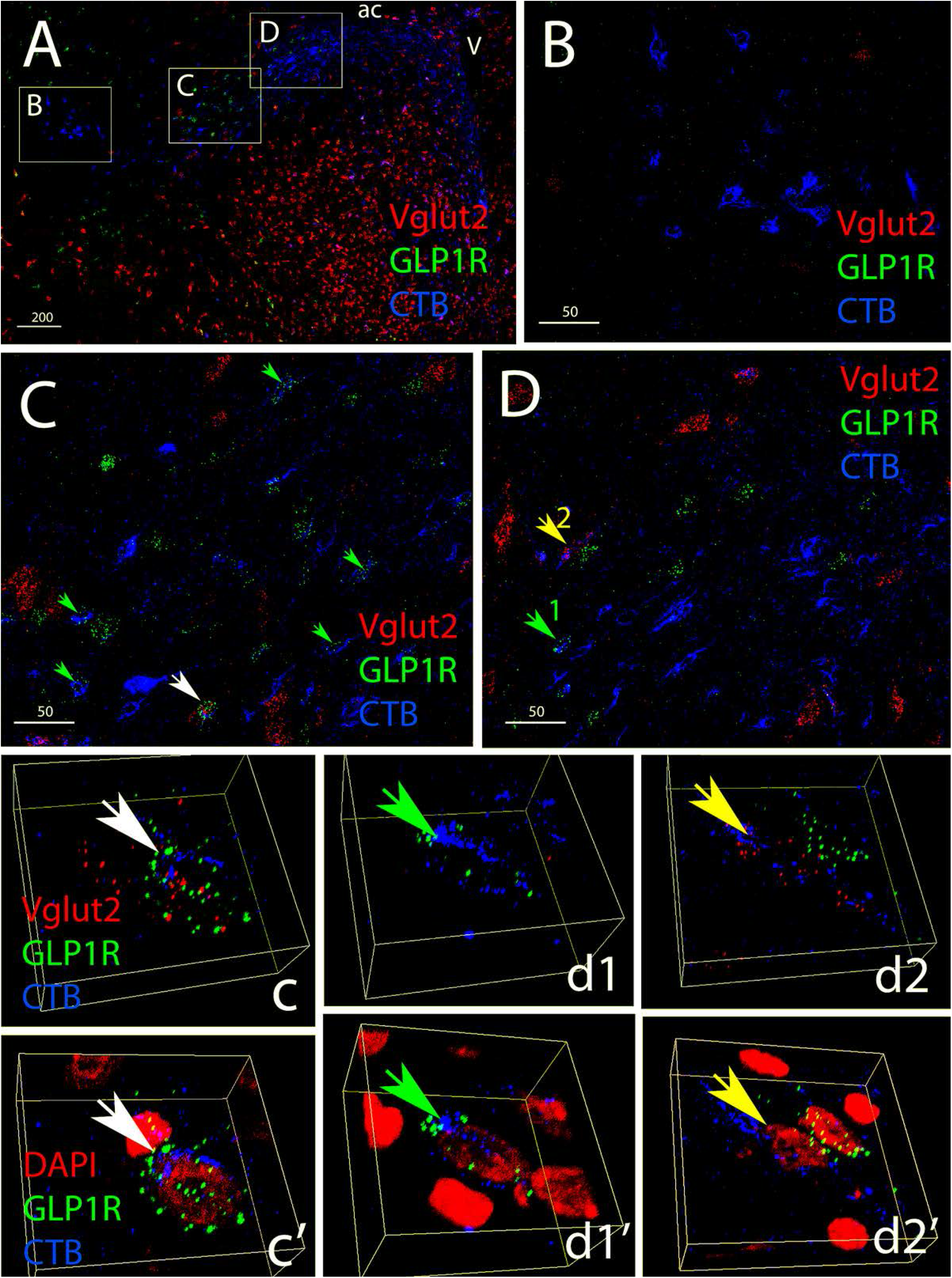
PVN-projecting neurons in the ventral alBST do not express Vglut2; PVN-projecting neurons in the anteromedial subnucleus rarely express GLP1R mRNA. **(A)** Low magnification Z-max projection image of a coronal section through the caudal portion of the left ventral alBST, approximately 0.5 mm caudal to bregma. Dorsal is towards the top, medial is towards the right. CTB labeling (blue) is imaged together with GLP1R mRNA (green) and Vglut2 mRNA (red). The boxed regions indicated by the labels B, C, and D (from lateral to medial) are shown at higher magnification in the correspondingly labeled panels. More medially (closer to the third ventricle), CTB-labeled neurons occupy the median preoptic and preoptic periventricular nuclei of the hypothalamus. ac, anterior commissure; V, third ventricle. Scale bar, 200 microns. **(B)** High magnification Z-max projection image from the boxed sub-region labeled B in panel A, corresponding to the location of the ventral anterolateral and rhomboid subnuclei of the BST. CTB-labeled neurons in this region do not express mRNA for Vglut2, although a few Vglut2-expressing cells are present. Sparse GLP1R mRNA expression is not colocalized with CTB. **(C)** High magnification Z-max projection image from the boxed sub-region labeled C in panel A, corresponding to the location of the anteromedial subnucleus of the BST. None of the CTB-labeled neurons in this region express Vglut2 mRNA, but several express GLP1R mRNA (confirmed examples of colocalization are indicated by arrowheads). The larger white arrowhead points out a CTB-labeled, GLP1R+ neuron located adjacent to a different Vglut2+ cell. These cells are shown at higher magnification in panel c, where 3D-MIP demonstrates colocalization of CTB (blue) and GLP1R mRNA (green) imaged along with Vglut2 (red). In c’, CTB and GLP1R mRNA is imaged along with nuclear DAPI (red). Scale bar, 50 microns. **(D)** High magnification Z-max projection image from the boxed sub-region labeled D in panel A, corresponding to the location of the dorsomedial subnucleus of the BST; a lateral portion of the hypothalamic median preoptic nucleus also is included in the upper (more dorsal) region of this field of view. Most CTB-positive neurons in this region express neither Vglut2 nor GLP1R mRNA; exceptions include the two neurons indicated by arrowheads. The green arrowhead (1) points out a CTB-labeled neuron that expresses GLP1R mRNA [shown in a higher magnification 3D-MIP in d1 (CTB plus GLP1R), with red DAPI nuclear labeling added in d1’]. The yellow arrowhead (2) points out a CTB-labeled neuron that expresses Vglut2 but not GLP1R mRNA [shown in a higher magnification 3D-MIP in d2 (CTB plus Vglut2 mRNA are colocalized in the same cell, whereas an adjacent cell expresses GLP1R but is not CTB-positive; red DAPI nuclear labeling is included with CTB and GLP1R labeling in d2’].

#### PVN-projecting neurons in the hypothalamic PS and ventral alBST are predominantly or exclusively GABAergic, and very few express GLP1R

CTB immunofluorescence was combined with fluorescent RNAscope *in situ* hybridization to colocalize retrograde labeling with GLP1R mRNA and Vgat mRNA, the latter to identify GABAergic neurons. Alternate sections from the same animals were processed to colocalize CTB, GLP1R mRNA, and Vglut2 mRNA, the latter to identify glutamatergic neurons. In each section analyzed, retrogradely-labeled (CTB-positive) neurons in different subnuclei of the ventral alBST and adjacent regions were inspected at low and high magnification to discern c0-expression of mRNA for GLP1R and either Vgat or Vglut2 mRNA.

Essentially all CTB-positive, PVN-projecting neurons within the fusiform, anterolateral, and anteromedial subnuclei of the ventral alBST and within the adjacent hypothalamic PS were GABAergic, based on visible labeling for Vgat mRNA (**Fig. 6**) and/or lack of Vglut2 mRNA labeling (**Fig. 7**). Surprisingly, and with rare exception, CTB-labeled neurons within the PS nucleus and ventral alBST did not express mRNA for GLP1R, and neurons within these regions that did express GLP1R mRNA were not tracer-labeled. Most of these GLP1 R-expressing neurons co-expressed Vgat mRNA (**Fig. 6**) rather than Vglut2 mRNA (**Fig. 7**). A thorough search yielded only rare examples (n=3) of CTB-labeled neurons within the ventral alBST/PS that appeared to express GLP1R mRNA. These three neurons did not co-express Vglut2 mRNA (suggesting a GABAergic phenotype), and were located in regions corresponding to the anteromedial and dorsomedial subnuclei of the ventral alBST (**Fig. 7**).

#### GLP1R is expressed by a subset of GABAergic neurons in the oval subnucleus of the dorsal alBST that are retrogradely labeled from the ventral alBST/PS

Consistent with previous reports (Dong et al., 2001)(Shin et al., 2008)(Bienkowski and Rinaman, 2013), iontophoretic delivery of CTB into the ventral alBST/PS produced retrograde labeling that was concentrated within the oval subnucleus of the dorsal alBST at the same rostro-caudal level (**Fig. 8**). All CTB-positive neurons within the oval subnucleus were GABAergic, based on their expression of Vgat mRNA (**Fig. 8**). A subset of these tracer-labeled neurons expressed mRNA for GLP1R (**Fig. 8**).

**Figure 8.**
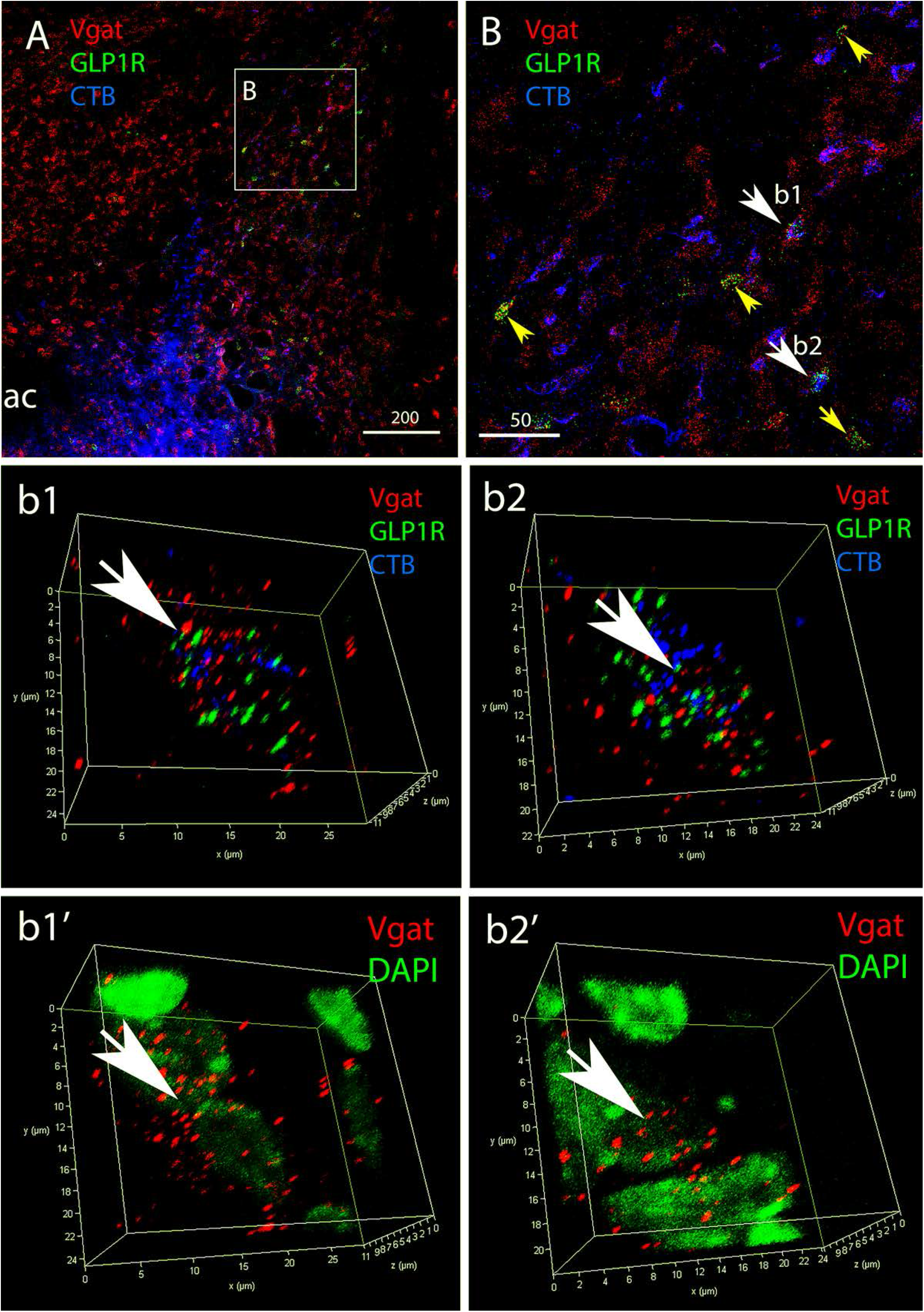
GABAergic neurons in the oval subnucleus of the dorsal alBST project into the ventral alBST, and a subset of these projection neurons express GLP1R mRNA. **(A)** Z-max projection image of a coronal section through the caudal portion of the right alBST, approximately 0.15 mm caudal to bregma. Dorsal is towards the top, medial is towards the left. The CTB iontophoretic tracer delivery site (lower part of image, adjacent to the ac) is centered in a region corresponding to the location of the ventral anteromedial, fusiform, and anterolateral subnuclei of the BST. CTB labeling (blue) is visible both within the injection site and retrogradely transported to the dorsal alBST. CTB labeling is imaged together with mRNA for Vgat (red) and GLP1R (green). The boxed region labeled B corresponds to the oval subnucleus of the dorsal alBST, shown at higher magnification in the correspondingly labeled panel. ac, anterior commissure. Scale bar, 200 microns. **(B)** Higher magnification Z-max projection image from the boxed sub-region labeled B in panel A, corresponding to the oval subnucleus of the dorsal alBST. All CTB-positive neurons (blue) express Vgat mRNA (red), and several also express GLP1R mRNA (green). In addition, many non-tracer-labeled neurons express Vgat mRNA, and some of these co-express GLP1R mRNA (indicated by small yellow arrowheads). The larger white arrowheads labeled b1 and b2 point out two triple-labeled neurons (i.e., positive for CTB, Vgat mRNA, and GLP1R mRNA). These cells are shown at higher magnification in 3D-MIP images in panels b1 and b2. For each cell, colocalization of all three labels is demonstrated in b1’ and b2’, which depict Vgat mRNA (red) and nuclear DAPI label (green).

## Discussion

We previously reported that ventral alBST neurons that express GLP1R are predominantly GABAergic in rats, and that virally-mediated “knock-down” of GLP1R expression in the alBST prolongs the plasma cort response to acute stress (Zheng et al., 2019). Based on evidence that a GABAergic projection pathway from ventral alBST to PVN serves to limit stress-induced activation of the HPA axis in rats (Radley et al., 2009)(Johnson et al., 2016), we hypothesized that the GABAergic neurons comprising this projection pathway are directly stimulated by GLP1. Results from the present study are consistent with the functional component of our hypothesis, but they challenge a specific detail. As predicted, pharmacological stimulation of GLP1R’s in *ex vivo* slices through the rat alBST promoted depolarization of identified PVN-projecting GABAergic neurons. However, with rare exception, PVN-projecting neurons within the ventral alBST did not express GLP1R mRNA, consistent with electrophysiological data indicating that the net excitatory effect of GLP1 signaling on these PVN-projecting neurons is indirect, and due to circuit-mediated effects within the *ex vivo* slice that increased excitatory synaptic inputs and decreased inhibitory synaptic inputs to the recorded neurons. We do not believe that retrograde labeling with CTB interfered with detection of GLP1R mRNA expression, because many examples of CTB-positive, GLP1R mRNA-expressing neurons were observed within the oval subnucleus of the dorsal alBST after tracer injection into the ventral alBST.

We acknowledge that whole-cell patch recordings in tracer-labeled neurons likely included cells within the hypothalamic PS, positioned immediately adjacent to the ventral alBST. The PS is not considered a component of the BST, because it does not receive axonal input from the amygdala (Ju and Swanson, 1989). However, the cytoarchitectural, connectional, and neurochemical properties of PS neurons are otherwise quite similar to those of neurons within the ventral alBST (Simerly and Swanson, 1988)(Thompson, 2003)(Dong and Swanson, 2006b). In this regard, results from the present study confirmed that virtually all PVN-projecting neurons within the ventral alBST and PS express Vgat mRNA, identifying them as GABAergic. This finding is consistent with previous reports that the majority of neurons within the ventral alBST and PS in rats are GABAergic (Sun and Cassell, 1993). Not surprisingly, virtually all alBST/PS neurons that express mRNA for GLP1R were seen to co-express Vgat mRNA. This finding is consistent with results from our previous study demonstrating co-expression of mRNA’s for GLP1R and a different GABAergic marker (i.e., Gad1) in the rat alBST (Zheng et al., 2019).

Data obtained from whole-cell patch recordings in *ex vivo* slices indicated that the intrinsic membrane properties of tracer-labeled (i.e., PVN-projecting) alBST/PS neurons closely resembled the membrane properties of nearby neurons that were not tracer-labeled (**Table 1**), and were consistent with previous reports describing electrophysiological properties of alBST neurons in rats and mice (Egli & Winder, 2003)(Hammack et al., 2007)(Rodríguez-Sierra et al., 2013)(Williams et al., 2018). As noted above, the depolarizing effect of Ex-4 on tracer-labeled neurons within the ventral alBST/PS appeared to be due to a reduction in inhibitory (IPSP) inputs coupled with an increase in excitatory (EPSP) inputs. Under conditions of synaptic blockade, Ex-4 produced no discernable effects on tracer-labeled neurons. These electrophysiological data suggested that PVN-projecting neurons within the ventral alBST/PS lack functional GLP1R’s, but that they receive synaptic input from neurons contained within the coronal *ex vivo* slice that are themselves directly or indirectly responsive to GLP1. Sources of such synaptic input likely include other local neurons that express GLP1R mRNA, although patch recordings made in non-tracer-labeled neurons located in close proximity to PVN-projecting neurons failed to identify any that responded (either directly or indirectly) to bath application of Ex-4. It is possible that increased sampling of non-labeled neurons would have revealed some that responded to Ex-4.

An additional projection pathway that was likely preserved within coronal *ex vivo* slices arises from the oval subnucleus of the dorsal alBST, which provides robust direct input to the subjacent ventral alBST at the same rostrocaudal level (Dong et al., 2001)(Shin et al., 2008). Noting that some neurons within the oval subnucleus express GLP1R mRNA, we combined *in situ* hybridization with retrograde labeling to reveal that neurons within the oval subnucleus that project to the ventral alBST are GABAergic, and that a subset of these neurons express GLP1R mRNA. In mice, activation of oval BST neurons leads to *inhibition* of postsynaptic ventral alBST neurons that project to the lateral hypothalamic area (Wang et al., 2019); this population of ventral alBST neurons may be distinct from those that project to the PVN. Our finding that bath application of Ex-4 promotes indirect, circuit-mediated *depolarization* of PVN-projecting alBST/PS neurons suggests that potential GLP1R-mediated effects on GABAergic inputs from the oval subnucleus contribute to disinhibition of PVN-projecting alBST/PS neurons. For example, GLP1R signaling could inhibit the GABAergic oval-to-ventral alBST projection, and/or could activate oval GABAergic neurons that synapse on another group of inhibitory interneurons that are presynaptic to PVN-projecting neurons in the ventral alBST. In this regard, GLP1R signaling in the mouse alBST directly depolarizes or hyperpolarizes different subpopulations of dorsal and ventral alBST neurons that apparently are intermixed (Williams et al., 2018), evidence that coupling to G_S_ protein (Fletcher et al., 2016) is not obligatory for all GLP1R-mediated neural responses [see also (Cork et al., 2015)(Ong et al., 2017)(Smith et al., 2019)].

To our knowledge, the current report is the first to examine GLP1R-mediated effects on the physiological properties of BST neurons with identified axonal projections. Previous studies conducted in rats and mice have reported a diversity of modulatory effects mediated by GLP1R signaling across different brain regions. For example, GLP1R signaling within the rat dorsomedial hypothalamus (DMH) appears to activate local GABAergic interneurons (Lee et al., 2018). Conversely, GLP1R signaling appears to exert a net inhibitory effect on GABAergic neurons within the mouse dorsolateral septum, where neurons become more excitable after genetic ablation of GLP1R (Harasta et al., 2015). Other studies have reported that GLP1R signaling reduces the excitability of neurons within the paraventricular thalamic nucleus that project to the nucleus accumbens, and that this effect is mediated in part via suppression of excitatory synaptic drive (Ong et al., 2017). On the other hand, GLP1R signaling enhances membrane trafficking of glutamate AMPA receptor subunits to thereby increase postsynaptic excitatory synaptic transmission within mouse PVN neurons (Liu et al., 2017). The latter finding suggests a potential mechanism underlying the increased excitatory postsynaptic currents that we recorded in ventral alBST neurons that project to the PVN; however, and in contrast to our current findings in rat alBST neurons, inhibitory postsynaptic currents were not altered by GLP1R signaling in mouse PVN neurons (Liu et al., 2017). The collective data support the view that GLP1R-mediated neural responses display a high degree of heterogeneity across different brain regions. Additional complexity arises from heterogeneity among neurons within each brain region; for example, ventral alBST/PS neurons that project to the PVN in rats are distinct from an intermingled population of midbrain-projecting neurons (Johnson et al., 2019), and it is possible that GLP1R signaling differentially impacts these distinct projections.

In summary, results from the present study provide evidence that GLP1R is not expressed by PVN-projecting neurons within the ventral alBST in rats. However, GLP1R-mediated effects on presynaptic inputs to these GABAergic neurons consistently promoted their depolarization. Considered together with results from previous studies (Radley et al., 2009)(Johnson et al., 2016)(Zheng et al., 2019), our findings reveal a potential GLP1R-mediated mechanism through which the ventral alBST exerts inhibitory control over the endocrine HPA axis. Additional research is necessary to unravel the potentially complex neural circuitry through which GLP1 neurons in the caudal brainstem engage GLP1R signaling pathways to exert this modulatory control, and the physiological conditions under which these circuits are engaged.

## Acknowledgments

We thank Alissa DePiro for assistance with confocal reconstructions.

